# Pathological C-terminal phosphomimetic substitutions alter the mechanism of liquid-liquid phase separation of TDP-43 low complexity domain

**DOI:** 10.1101/2024.03.21.586202

**Authors:** Raza Haider, Brandon Shipley, Krystyna Surewicz, Michael Hinczewski, Witold K Surewicz

## Abstract

C-terminally phosphorylated TAR DNA-binding protein of 43 kDa (TDP-43) marks the proteinaceous inclusions that characterize a number of age-related neurodegenerative diseases, including amyotrophic lateral sclerosis, frontotemporal lobar degeneration and Alzheimer’s disease. TDP-43 phosphorylation at S403/S404, and especially at S409/S410, is in fact accepted as a biomarker of proteinopathy. These residues are located within the low complexity domain (LCD), which also drives the protein’s liquid-liquid phase separation (LLPS). The impact of phosphorylation at these LCD sites on phase separation of the protein is a topic of great interest, as these post-translational modifications and LLPS are both implicated in proteinopathies. Here, we employed a combination of experimental and simulation-based approaches to explore this question on a phosphomimetic model of the TDP-43 LCD. Our turbidity and fluorescence microscopy data show that Ser-to-Asp substitutions at residues S403, S404, S409 and S410 alter the LLPS behavior of TDP-43 LCD. In particular, in contrast to the unmodified protein, the phosphomimetic variants display a biphasic dependence on salt concentration. Through coarse-grained modeling, we find that this biphasic salt dependence is derived from an altered mechanism of phase separation, in which LLPS-driving short-range intermolecular hydrophobic interactions are modulated by long-range attractive electrostatic interactions. Overall, this *in vitro* and *in silico* study provides a physiochemical foundation for understanding the impact of pathologically-relevant C-terminal phosphorylation on the LLPS of the TDP-43 in a more complex cellular environment.

**Statement of Significance:** Proteinaceous inclusions composed of phosphorylated, C-terminal TDP-43 fragments have long been recognized as hallmarks of several neurodegenerative diseases, in particular amyotrophic lateral sclerosis and frontotemporal dementia. A rapidly growing number of studies indicate that these proteinopathies may be closely related to liquid-liquid phased separation (LLPS) of TDP-43, but the impact of phosphorylation on TDP-43 LLPS remains largely unexplored. In this study we used a combination of experimental methods and coarse-grained simulations to ascertain, in mechanistic terms, how phosphorylation at pathologically-critical C-terminal sites impacts liquid-liquid phase separation of the low complexity domain of TDP-43. Our results broaden our understanding of the mechanisms driving pathogenic process in these neurodegenerative diseases.

## Introduction

TAR DNA-binding protein of 43 kDa (TDP-43) is a nucleocytoplasmic nucleic-acid binding protein. Proteinaceous inclusions containing this protein typify a multitude of age-related neurodegenerative diseases, including amyotrophic lateral sclerosis, frontotemporal lobar degeneration, Alzheimer’s disease, limbic predominant age-related TDP-43 encephalopathy and cerebral age-related TDP-43 with sclerosis (Neumann et al. 2006; Arai et al. 2006; Prasad et al. 2019; de Boer et al. 2020; Meneses et al. 2021). Previous studies have established that post-translational modifications such as N-terminal truncation, ubiquitination and phosphorylation may play an important role in the protein’s misfolding and aggregation (Neumann et al. 2006; Arai et al. 2006; Neumann et al. 2009; Nonaka et al. 2009). Phosphorylation appears to be of particular importance in this regard, as pathognomonic proteinaceous inclusions are marked by hyperphosphorylated TDP-43 (Neumann et al. 2009; Hasegawa et al. 2008). Even though many TDP-43 phosphorylation sites have been identified (Eck, Kraemer, and Liachko 2021), phosphorylation of residues in the intrinsically-disordered low complexity domain (LCD; res. ∼267-414) appear to be especially strongly linked to disease (Eck, Kraemer, and Liachko 2021; Hasegawa et al. 2008; Neumann et al. 2009; Neumann et al. 2021). Of these, four at the very C-terminal end of the protein (i.e., S403, S404, S409, and S410) are the best established, to the point that the presence of TDP-43 phosphorylated at these sites is an accepted biomarker of disease (Eck, Kraemer, and Liachko 2021; Hasegawa et al. 2008; Neumann et al. 2009).

TDP-43 is stratified into four canonical domains – the N-terminal domain, which is involved in oligomerization of the protein, two RNA recognition motifs, and the C-terminal, intrinsically-disordered LCD (Prasad et al. 2019; François-Moutal et al. 2019). In the context of neurodegenerative diseases, the LCD has received particular attention, as patient-derived inclusions are enriched in C-terminal fragments containing this domain (Neumann et al. 2006; Arai et al. 2006; Igaz et al. 2008; Arseni et al. 2022). The LCD also appears to be critical for protein aggregation in experimental models (Johnson et al. 2009; Furukawa et al. 2011; Yang et al. 2010), and houses most of the disease-related mutations of TDP-43 (Prasad et al. 2019; Buratti 2015; Eck, Kraemer, and Liachko 2021). Furthermore, recent studies have revealed that the LCD is essential for liquid-liquid phase separation (LLPS) of TDP-43 (Conicella et al. 2016; Schmidt and Rohatgi 2016; Carey and Guo 2022), the phenomenon through which protein condenses into reversible, liquid-like droplets (Banani et al. 2017; Shin and Brangwynne 2017; Alberti and Dormann 2019). This is of particular interest since growing evidence points to a major role of LLPS in protein aggregation and the pathogenesis of age-related neurodegenerative diseases (Alberti and Dormann 2019; Wang et al. 2021; Babinchak and Surewicz 2020; Zbinden et al. 2020; Haider, Boyko, and Surewicz 2023).

LLPS of the TDP-43 LCD is believed to be driven by multiple intermolecular forces working in concert. Among those, hydrophobic interactions between multiple non-polar residues (Schmidt, Barreau, and Rohatgi 2019; Babinchak et al. 2019; Babinchak et al. 2020; Li, Chen, et al. 2018) as well interactions involving aromatic side chains (e.g., cation-π, π-π stacking, methionine-phenylalanine) (Schmidt, Barreau, and Rohatgi 2019; Li, Chiang, et al. 2018; Mohanty et al. 2023) appear to be especially important, since addition of compounds that disrupt hydrophobic forces and removal of key Trp and Phe residues severely inhibit LLPS. Of equal importance appears to be the transiently α-helical region, a segment within the LCD encompassing residues ∼320 – 340. Indeed deletion of this segment or helix-breaking mutations within it lead to the abolition of protein LLPS (Conicella et al. 2016; Schmidt and Rohatgi 2016; Jiang et al. 2013; Li, Chiang, et al. 2018; Schmidt, Barreau, and Rohatgi 2019; Li, Chen, et al. 2018; Haider et al. 2024). On the other hand, electrostatic interactions between the relatively few charged residues natively present within the LCD (concentrated mainly within the N-terminal part of the protein) result in repulsive intermolecular electrostatic forces between monomers, leading to inhibition of LLPS at low ionic strength (Li, Chen, et al. 2018; Babinchak et al. 2019).

The balance of these forces is likely to be substantially impacted by specific post-translational modifications, affecting LLPS of modified protein. However, little information is available in this regard, especially for protein phosphorylated at pathologically critical sites within the C-terminal part of the LCD. Here we attempt to bridge this gap using a phosphomimetic approach in which Ser residues at position 403, 404, 409, and 410 have been replaced with Asp. Our experimental data reveal that these phosphomimetic substitutions alter the LLPS behavior of TDP-43 LCD, especially with regard to its dependence on ionic strength. Coarse-grained simulations provide insight into the mechanism of this altered behavior, suggesting that it is due to a different balance of hydrophobic and electrostatic forces in the non-modified and phosphomimetic protein.

## Results

### C-terminally phosphomimetic substitutions confer a biphasic salt dependence to TDP-43 LCD LLPS

To investigate how phosphorylation of C-terminal Ser residues (i.e., S403, S404, S409, and S410) in TDP-43 LCD affects protein LLPS, we generated phosphomimetic variants of the LCD, with varying numbers of Ser-to-Asp substitutions to model phosphorylation at two (S403D/S404D, referred to as 2-PM TDP-43) or four (S403D/S404D/S409D/S410D, referred to as 4-PM TDP-43) of the aforementioned C-terminal residues (**Fig. 1A**). We then measured the turbidity (the optical density at 600 nm) of these variants at 20 µM concentration and physiological pH as a function of NaCl concentration to see what effect the C-terminal phosphomimetic substitutions had on TDP-43 LCD LLPS. Wild-type (WT) TDP-43 LCD LLPS showed a direct dependence on ionic strength, with increasing salt concentration promoting phase separation monotonically, in agreement with other studies (Conicella et al. 2016; Li, Chiang, et al. 2018; Babinchak et al. 2019). The LLPS of the phosphomimetic TDP-43 LCD proteins, in contrast, were characterized by a biphasic salt dependence. At low-to-moderate NaCl concentrations, 2-PM TDP-43 LCD and 4-PM TDP-43 LCD showed diminished LLPS in response to increasing ionic strength, opposite to the behavior of WT protein. However, at moderate-to-high NaCl concentrations, 2-PM TDP-43 LCD and 4-PM TDP-43 LCD showed increased LLPS in response to increasing ionic strength; this switch occurred at a different salt concentration for each phosphomimetic variant (**Figs. 1B-C**). Furthermore, the droplets formed by 4-PM TDP-43 LCD were also much smaller than those formed by the other proteins.

**Figure 1.**
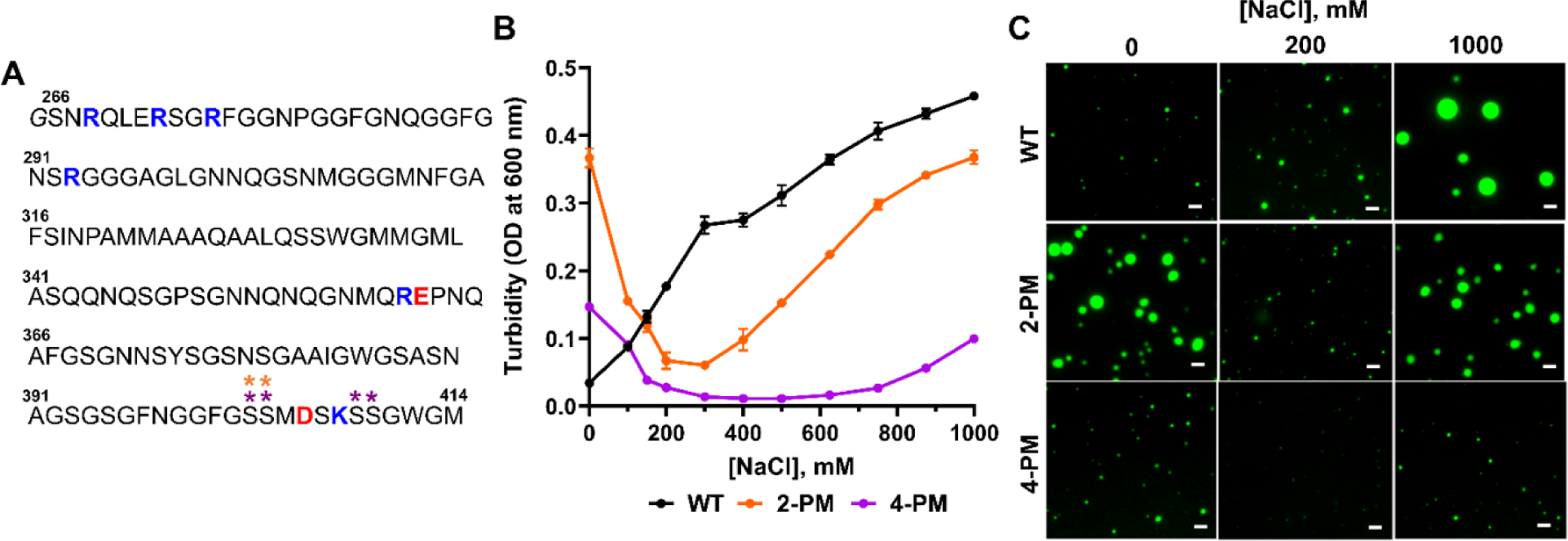
The LLPS of C-terminal phosphomimetic TDP-43 LCD variants shows a non-monotonic salt response. *A*, Sequence of the TDP-43 low complexity domain construct used in this study. An extra N-terminal glycine from the purification process is *italicized*, and natively positively- and negatively-charged residues are marked in blue and red, respectively. The residues that were phosphomimetically-substituted to form the 2-PM variant are marked with orange asterisks, and those of the 4-PM variant are marked with purple asterisks. *B*, Turbidity plots of TDP-43 LCD variants (20 µM in each case) as a function of ionic strength. Error bars represent SD (n=3). *C*, Representative fluorescence microscopy images of TDP-43 LCD variants at 20 µM protein. Proteins were labeled with Alexa Fluor 488, and the ratio of labeled-to-unlabeled protein was 1:20 for each protein. Scale bar, 3 µm. Experiments were performed in 20 mM potassium phosphate buffer (pH 7.4), and data/images were collected ∼20 minutes after sample preparation. OD, optical density.

In order to more fully document these surprising results, saturation concentrations (*csat*) were calculated and phase diagrams were constructed for all of the TDP-43 LCD variants (**Figs. 2A-B**). The phase diagrams substantiated the non-monotonic salt dependences of the phosphomimetic proteins’ LLPS, illustrating that the concentrations of protein at which phase separation was triggered at first increased with increasing ionic strength, and then decreased as ionic strength was increased further (**Fig. 2B**). This behavior was reflected in the trends of the saturation concentrations (**Fig. 2A**). The salt concentration range in which the LLPS of a given phosphomimetic variant had an inverse dependence on ionic strength was denoted its “low salt regime,” and the range in which it showed a direct dependence on ionic strength was denoted its “high salt regime.”

**Figure 2.**
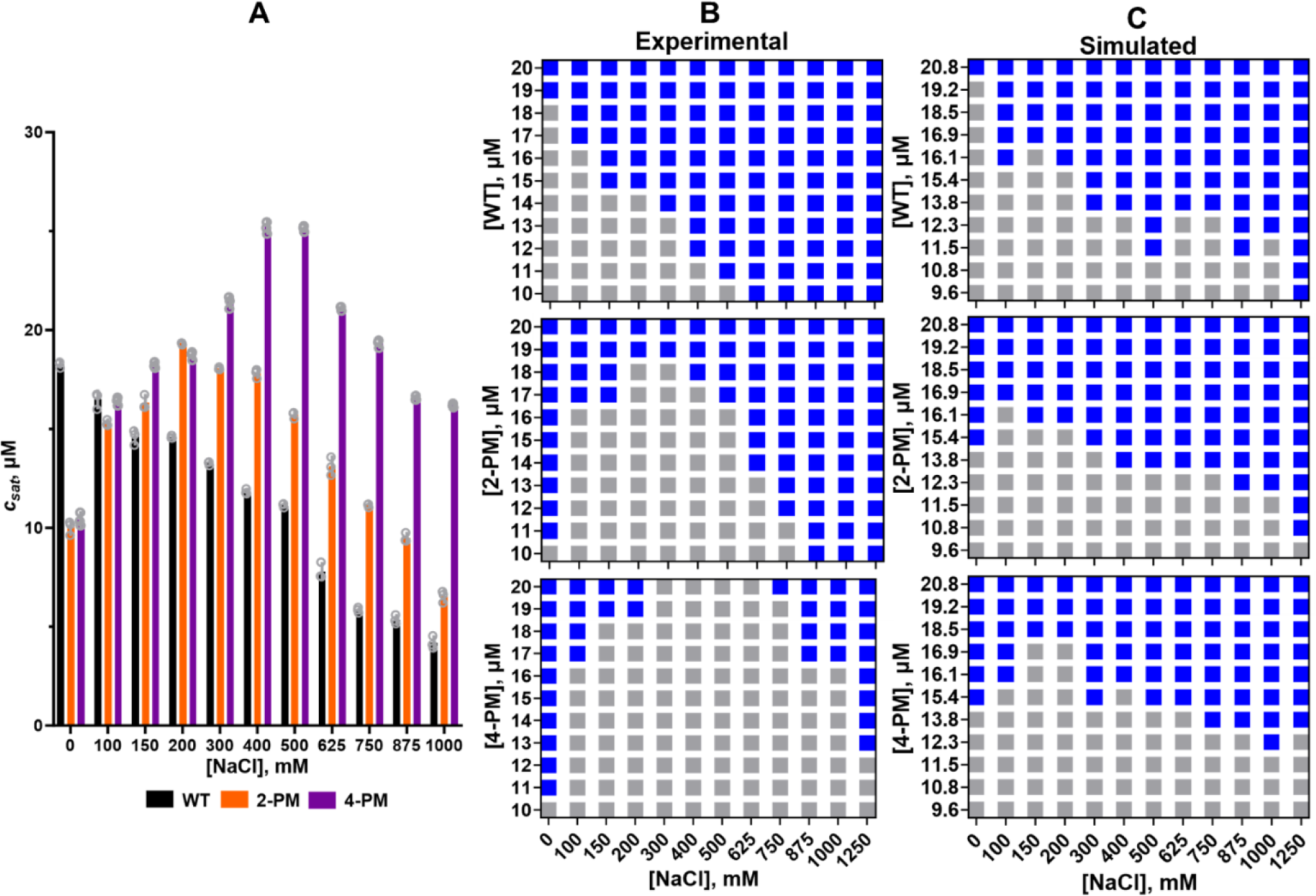
Phase diagrams and saturation concentrations of TDP-43 LCD variants. *A*, Experimentally determined saturation concentrations for TDP-43 LCD variants. *B*, Experimentally determined phase diagrams. *C*, *In silico* phase diagrams. The experiments and the *in silico* simulations were performed in 20 mM potassium phosphate buffer (pH 7.4), at room temperature.

### Hydrophobic forces are predominantly responsible for TDP-43 LCD phase separation

A number of intermolecular forces have been shown to contribute to WT TDP-43 LCD LLPS. The LCD is enriched in hydrophobic amino acids, and hydrophobic interactions between these residues have a key role in driving the protein’s phase separation (Schmidt, Barreau, and Rohatgi 2019; Babinchak et al. 2019; Babinchak et al. 2020; Li, Chen, et al. 2018). A segment of the LCD spanning residues ∼320 – 340 that has high propensity to form a transient α-helix has been revealed to be especially important, as it mediates intermolecular helix-helix contacts without which LLPS is abrogated (Conicella et al. 2016; Schmidt and Rohatgi 2016; Jiang et al. 2013; Li, Chiang, et al. 2018; Schmidt, Barreau, and Rohatgi 2019; Li, Chen, et al. 2018; Haider et al. 2024). Additionally, there is evidence that repulsive intermolecular electrostatic interactions arising from the net +3 charge of the protein (concentrated mainly at its N-terminal end) are at play, inhibiting LLPS at low salt concentrations (Li, Chen, et al. 2018; Babinchak et al. 2019). The biphasic salt response observed for LLPS of the C-terminal phosphomimetic variants, however, suggested that electrostatic forces may drive condensation of these variants in the low salt regime, and hydrophobic forces take over in the high salt regime.

To investigate the roles of electrostatic and hydrophobic forces in the phase separation of C-terminally phosphomimetic TDP-43 LCD, we set up sequence-dependent coarse-grained simulations. Our methods were based on established techniques, which have previously been used to study LLPS of TDP-43 LCD (Gruijs da Silva et al. 2022), as well as other proteins such as the LCD of the RNA-binding protein FUS (Dignon et al. 2018; Monahan et al. 2017; Joseph et al. 2021), the DEAD-box helicase LAF-1 (Dignon et al. 2018), and the RNA-binding protein hnRNPA2 (Ryan et al. 2018). In this system, amino acids were represented as spheres with effective masses, charges, hydrophobicities and sizes taken from the literature (**Table S1**) (Dignon et al. 2018). A Yukawa screened Coulombic potential and a modified Lennard-Jones potential of the Ashbaugh Hatch functional form (Ashbaugh and Hatch 2008) were used to implement electrostatic and hydrophobic forces, respectively. Phase diagrams were then generated *in silico* in order to verify that the coarse-grained model captured the biphasic salt dependence the LLPS of the phosphomimetic variants displayed in our experiments *in vitro*. Despite some quantitative differences, these simulated phase diagrams did indeed recapitulate the non-monotonic salt dependences of the phosphomimetic proteins, as well as the monotonic salt dependence of the WT protein within a similar protein concentration range as observed in the experiments (**Fig. 2C**). Considering the unavoidable approximation used in coarse-grained simulations, such a good correspondence between experimental and simulated phase diagrams is quite remarkable, validating our *in silico* model. Of note, to achieve these results, the well depth (ε) of the modified Lennard-Jones potential was explicitly made to scale with ionic strength (∼3% increase from 0 – 1250 mM NaCl) so as to represent the kosmotropic nature of Na+ ions and the inherent salt dependence of hydrophobic forces (Moelbert, Normand, and De Los Rios 2004). In the absence of this perturbation, simulated phase separation diagrams notably did not recapitulate the qualitative trends of their experimental counterparts (**Fig. S1**).

Using the *in silico* data, we next compared how the total intermolecular electrostatic force changed for each TDP-43 variant at one selected protein concentration (20.8 µM) as the protein transitioned from the low to the high salt regime. Our analysis focused on the 20.8 µM simulations because at this protein concentration all three LCD variants phase separated at all salt concentrations tested *in silico* (**Fig. 2C**). The data showed that WT protein experienced a net repulsive electrostatic force at 0 mM NaCl that weakened as ionic strength was increased (**Fig 3A**), in line with expectations as WT TDP-43 LCD has a net positive charge. The phosphomimetic variants, in contrast, experienced a net attractive electrostatic force at 0 mM NaCl (which also weakened as ionic strength was increased) (**Figs. 3A and S2**), even though 2-PM TDP-43 and 4-PM TDP-43 also have net charges (of +1 and -1, respectively). This attractive electrostatic force likely arose from the polarized charge distribution of the phosphomimetic proteins. Unlike WT TDP-43, which has only a positively-charged N-terminal region, 2-PM TDP-43 and 4-PM TDP-43 have both the native N-terminal “positive pole” and a C-terminal “negative pole” (**Fig. 3B**), which could interact intermolecularly to generate the attractive electrostatic forces experienced by the phosphomimetic proteins in the low salt regime. These results at first suggested that the inverse salt dependence exhibited by the phosphomimetic TDP-43 variants could be due to attractive electrostatic forces driving their LLPS in their low salt regimes, and that the reversal of this relationship in the high salt regimes may be because the increased ionic strength screened out the electrostatic forces and allowed hydrophobic forces to take over.

**Figure 3.**
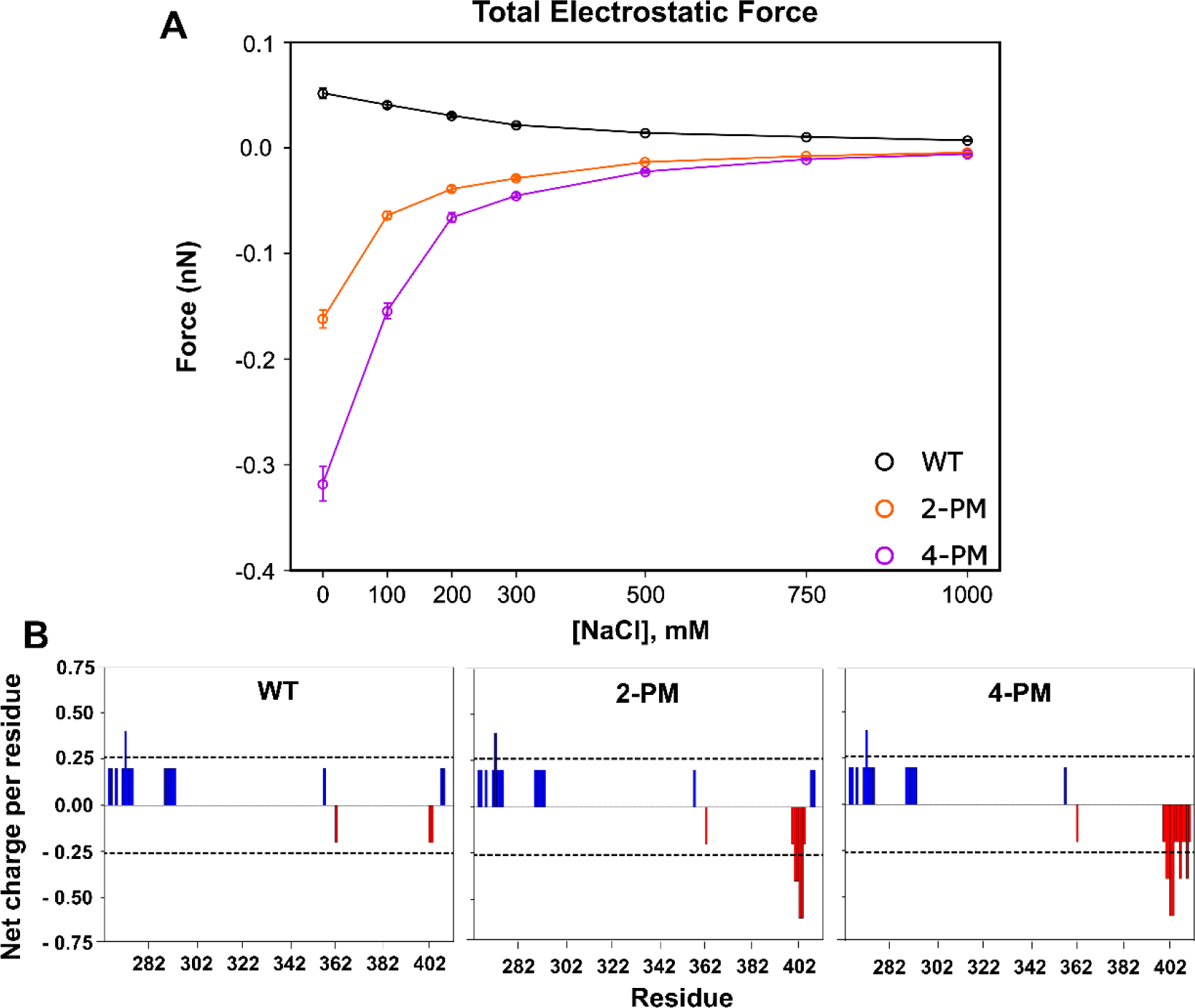
Total intermolecular electrostatic force generated by TDP-43 LCD interactions for each variant under the conditions of phase separation. *A,* Sum of total intermolecular electrostatic forces acting on TDP-43 LCD variants as a function of salt concentration, derived from simulations of 20.8 μM protein. *B,* Net charge per residue plots for TDP-43 LCD variants, generated using the CIDER tool (**REF**). Charge was calculated over a 5 residue window.

Examining the total intermolecular hydrophobic forces, however, cast some doubt regarding the importance of electrostatic forces in TDP-43 phase separation. Indeed, for each protein at every salt concentration surveyed, the total hydrophobic force exceeded the total electrostatic force by two orders of magnitude (**Figs. 3A and 4A**). Furthermore, at 0 mM NaCl, the total hydrophobic force experienced by the phosphomimetic TDP-43 variants was stronger than that experienced by WT protein (**Fig. 4A**). This analysis indicated that hydrophobic forces were the major driver of LLPS of not only WT TDP-43, but also of 2-PM TDP-43 and 4-PM TDP-43, in both the low salt and high salt regimes. To experimentally test this simulation-suggested possibility, we formed droplets of all protein variants studied in 0 mM NaCl, where electrostatic forces would be at their strongest, and exposed them to 1,6-hexanediol, an aliphatic alcohol that disrupts hydrophobic interactions (Kroschwald et al. 2015). 1,6-hexanediol did, in fact, dissolve these droplets (**Fig. 3B**), just as it did droplets formed at 1000 mM NaCl (**Fig. 3C**), supporting the notion that hydrophobic forces were the primary drivers of phase separation for all three proteins regardless of ionic strength.

**Figure 4.**
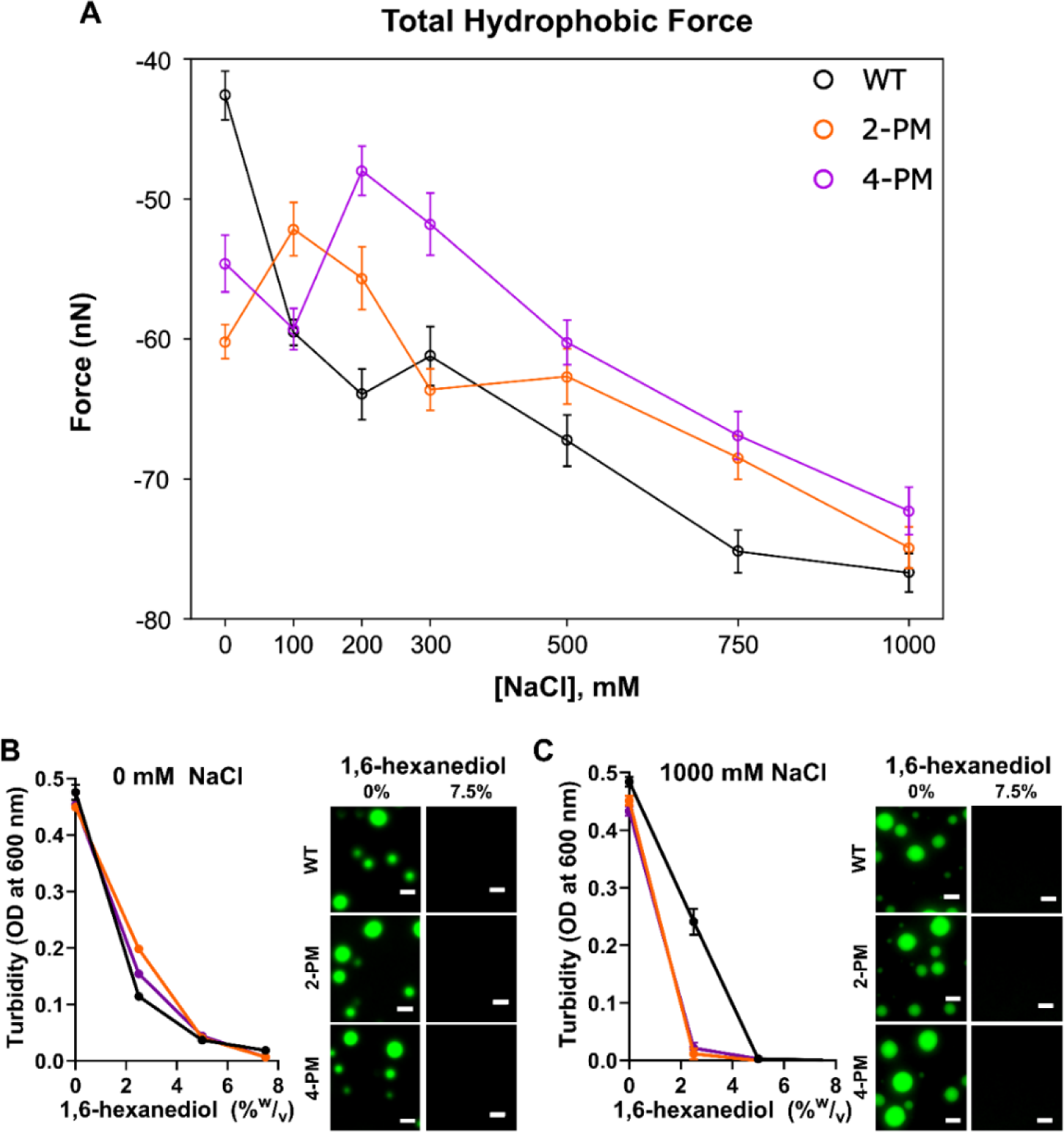
Hydrophobic forces dominate LLPS of WT TDP-43 LCD and phosphomimetic TDP-43 LCD both the low salt and high salt regimes. *A,* Sum of total intermolecular hydrophobic forces acting on TDP-43 LCD variants as a function of salt concentration, derived from simulations at a protein concentration of 20.8 μM. *B,* Sensitivity of TDP-43 droplets formed at 0 mM NaCl to 1,6-hexanediol, as shown by turbidity (*left*) and representative microscopy images (*right*). *C,* Sensitivity of TDP-43 droplets formed at 1000 mM NaCl to 1,6-hexanediol, as shown by turbidity (*left*) and representative microscopy images (*right*). Proteins were labeled with Alexa Fluor 488, and the ratio of labeled-to-unlabeled protein was 1:20 for each protein. Scale bar, 3 µm. Experiments were performed in 20 mM potassium phosphate buffer (pH 7.4), and data/images were collected ∼20 minutes after sample preparation. OD, optical density.

### Electrostatic forces tune the hydrophobic forces driving phosphomimetic TDP-43 LCD LLPS

Though inspecting the total intermolecular electrostatic and hydrophobic forces revealed that the latter were the major driver of phase separation of WT TDP-43 and phosphomimetic TDP-43 at all salt concentrations tested, this finding did not explain the inverse salt dependences of 2-PM TDP-43 LLPS and 4-PM TDP-43 LLPS in their low salt regimes. Due to the kosmotropic nature of Na+ ions (Moelbert, Normand, and De Los Rios 2004), NaCl enhances the hydrophobic effect, and would theoretically promote hydrophobic forces. The observation that NaCl inhibited LLPS of phosphomimetic TDP-43 in the low salt regime, however, suggested that electrostatic interactions still played a significant role. To explore this apparent conundrum, we constructed plots of the pairwise intermolecular electrostatic and hydrophobic forces occurring between every pair of residues of each protein at 0 mM NaCl (firmly in the low salt regime), 200 mM NaCl (near the transition point between the low and high salt regimes of 2-PM and 4-PM TDP-43) and 1000 mM NaCl (firmly in the high salt regime) (**Figs. 5, 6 and S1**). This analysis allowed us to understand how electrostatic and hydrophobic interactions between different regions of the proteins contributed to LLPS across the low and high salt regimes.

**Figure 5.**
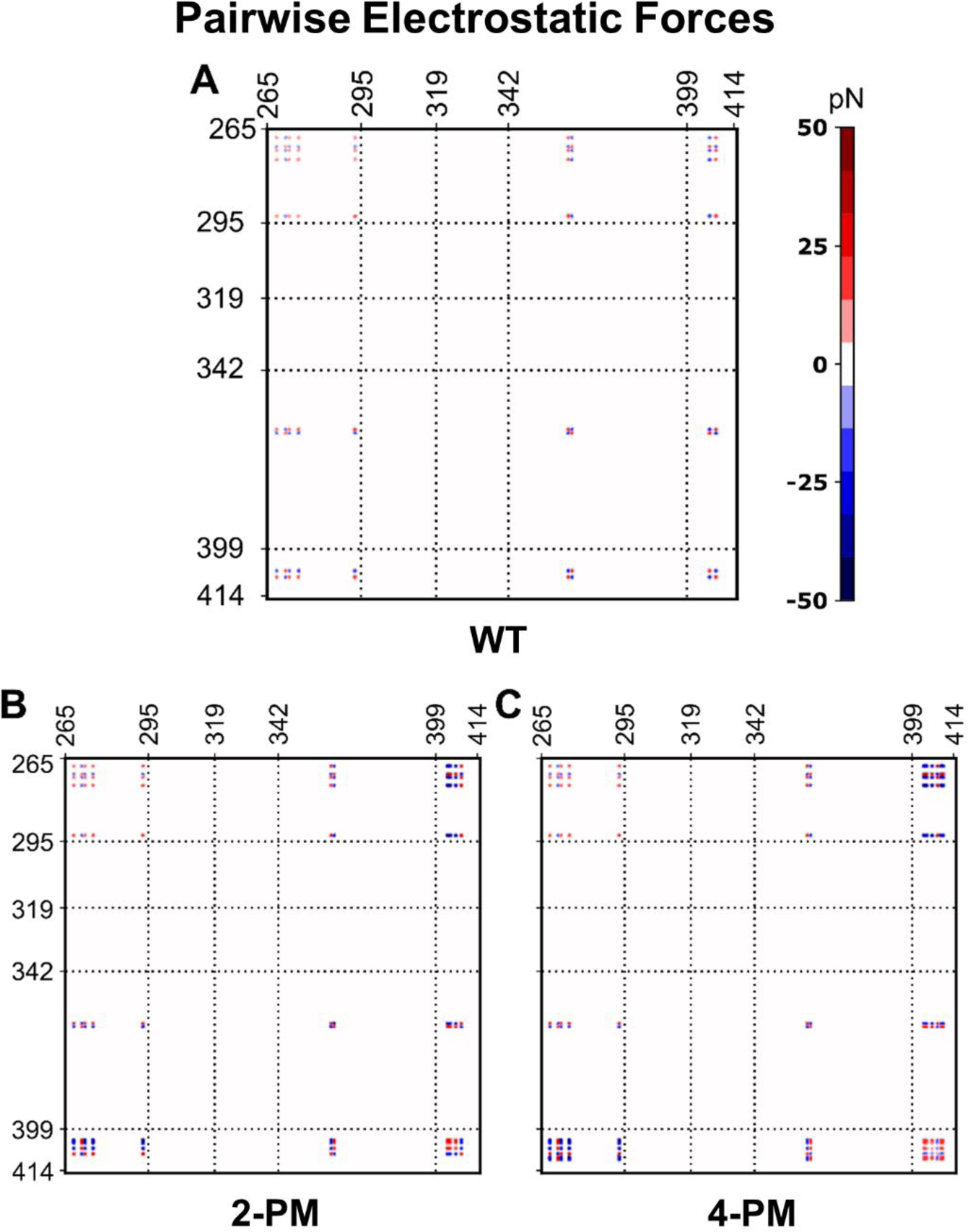
Intermolecular residue-residue pairwise electrostatic force plots of phase-separated TDP-43 LCD variants in the absence of salt. Pairwise intermolecular electrostatic forces of *A,* WT TDP-43 LCD, *B,* 2-PM TDP-43 LCD and *C,* 4-PM TDP-43 LCD. Data were generated from simulations at a protein concentration of 20.8 µM in a buffer containing no NaCl. Dashed reference lines mark the same residues in all three plots.

Interestingly, our simulations at 0 mN NaCl revealed that individual electrostatic forces were of the same order of magnitude as individual hydrophobic forces (**Figs. 5 and 6**). Thus, it appeared that the vastly larger magnitude of the total intermolecular hydrophobic force as compared to total intermolecular electrostatic force for each protein in the low salt regime (**Figs. 3A and 4A**) was largely due to the paucity of residues that were charged (8 – 12 residues) versus those able to interact hydrophobically (151 residues), rather than individual electrostatic forces being weaker than individual hydrophobic forces. Additionally, the pairwise electrostatic force data confirmed that the phosphomimetic proteins experienced net N-/N-terminal and net C-/C-terminal electrostatic repulsion, as well as net N-/C-terminal electrostatic attraction, at 0 mM NaCl, whereas WT protein experienced only N-/N-terminal repulsion at this ionic strength (**Fig. 5**). At 200 mM NaCl, these electrostatic interactions were still present, though weaker because of salt screening, and by 1000 mM NaCl, individual electrostatic forces were virtually non-existent (**Fig. S2**). These data thus suggested that the orientations of the phosphomimetic proteins within droplets differed from that of the WT protein in their low-salt regimes, but that they packed similarly to WT protein within droplets in their high salt regimes.

**Figure 6.**
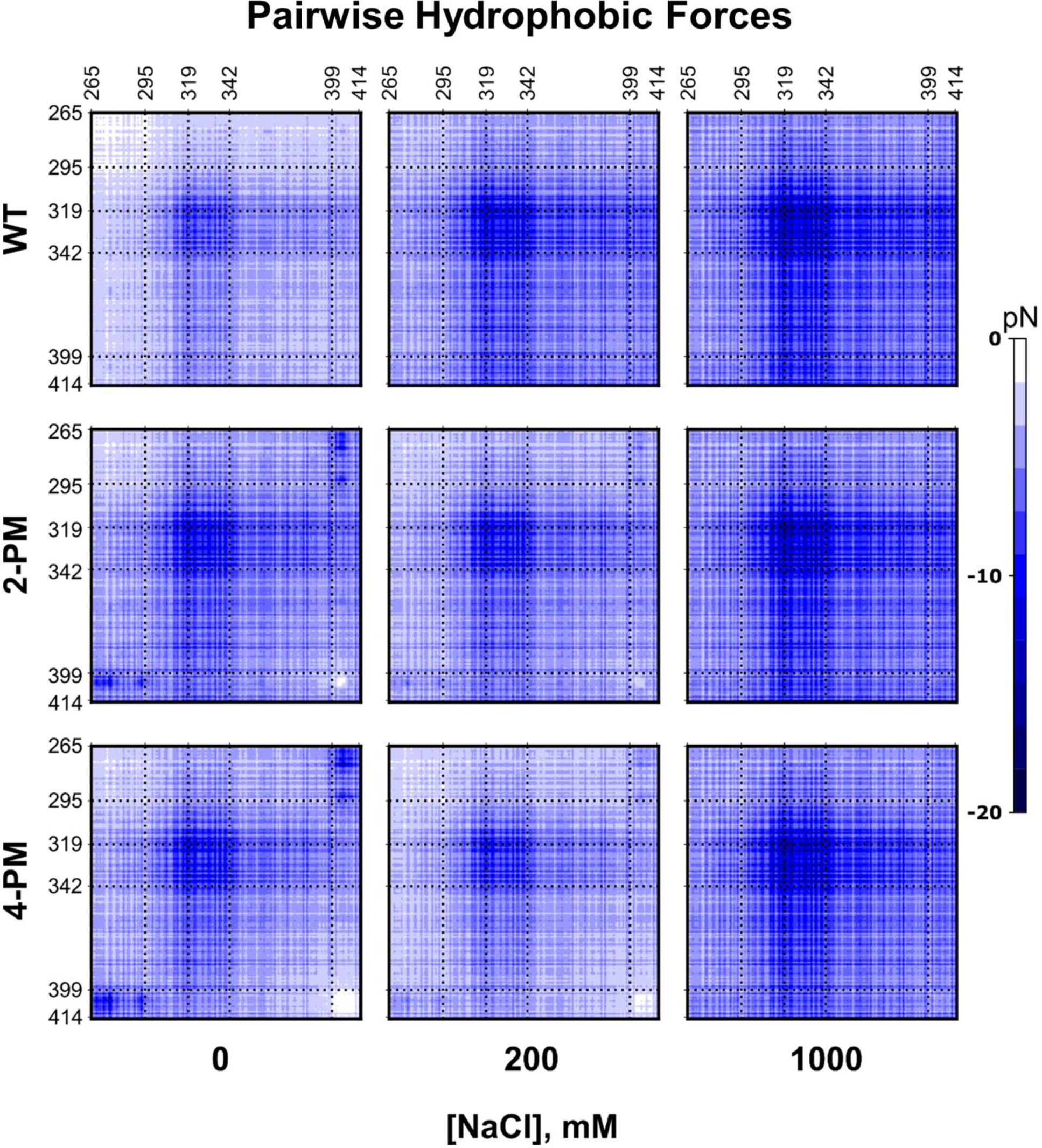
Intermolecular residue-residue pairwise hydrophobic force plots of phase-separated TDP-43 LCD variants. Pairwise intermolecular hydrophobic forces of WT TDP-43 LCD (*Top*), 2-PM TDP-43 LCD (*Middle*) and 4-PM TDP-43 LCD (*Bottom*) at various ionic strengths. Data were generated from simulations at a protein concentration of 20.8 µM. Dashed reference lines mark the same residues in all nine plots.

The apparent differences in the orientations of WT TDP-43 and phosphomimetic TDP-43 within droplets discussed above suggested that electrostatic interactions could tune the hydrophobic forces driving phase separation. Hydrophobic forces are exquisitely sensitive to distance, showing sharp exponential decay as the distance between interacting hydrophobic surfaces is enlarged. While ionically screened electrostatic forces also decay exponentially as the distance between interactions particles is increased, this decay is generally less prominent relative to hydrophobic forces at low salt concentrations (Israelachvili and Pashley 1982). Any electrostatic interaction that brings hydrophobic surfaces closer together would result in a stronger hydrophobic force between them. Conversely, any electrostatic interaction that pushes the surfaces farther apart would weaken said hydrophobic force. The pairwise hydrophobic force plots demonstrated this was indeed the case. At 0 mM NaCl, the pairwise hydrophobic forces from C-/C-terminal interactions were stronger for WT protein, which had no significant electrostatic forces between these regions, than for the phosphomimetic proteins, which had repulsive electrostatic forces between these regions. On the other hand, the pairwise hydrophobic forces from N-/C-terminal interactions were weaker for WT TDP-43 compared to those of 2-PM TDP-43 and 4-PM TDP-43, in accordance with the corresponding electrostatic forces being negligible for the former and attractive for the latter at this ionic strength.

The modulatory effect of electrostatic forces on the hydrophobic forces driving phosphomimetic TDP-43 LLPS was also evident as ionic strength was increased. For WT protein, as NaCl concentration was increased from 0 to 200 to 1000 mM, the pairwise intermolecular hydrophobic forces became stronger. For the 2-PM and 4-PM variants, in contrast, the pairwise intermolecular hydrophobic forces became weaker from 0 to 200 mM NaCl, but became stronger from 200 to 1000 mM NaCl. In fact, at 1000 mM NaCl, the phosphomimetic proteins’ plots were practically indistinguishable from that of the WT protein (**Fig. 6**).

Taken together, these results indicated a potential mechanism for the biphasic salt response of phosphomimetic TDP-43 LLPS. For WT protein, salt strengthened the hydrophobic forces mediating LLPS in a two-fold manner: (i) by enhancing the hydrophobic effect due to its kosmotropic nature and (ii) by screening repulsive N-/N-terminal electrostatic interactions. For the phosphomimetic variants, on the other hand, these two effects did not work in concert. While increasing NaCl concentration still enhanced the hydrophobic effect, the attractive N-/C-terminal electrostatic interactions that promoted hydrophobic forces driving LLPS were reduced. This led to overall weaker hydrophobic forces and inhibited LLPS for 2-PM TDP-43 LCD and 4-PM TDP-43 LCD as NaCl concentration was increased in the low salt regime. In the high salt regime, however, electrostatic forces were completely screened out; thus increasing ionic strength solely strengthened the hydrophobic forces (and thus LLPS), mirroring the behavior of WT protein.

### The transiently α-helical region of TDP-43 remains essential for LLPS of C-terminally phosphomimetic variants of TDP-43 LCD

The pairwise hydrophobic force plots for WT TDP-43, 2-PM TDP-43, and 4-PM TDP-43 all showed that the strongest hydrophobic forces experienced by the proteins in their phase separated states were generated by a stretch of amino acids interacting intermolecularly with itself (**Fig. 6**). This stretch, which encompassed residues ∼317-342, roughly corresponded to the transiently α-helical region of TDP-43. Thus, it appeared that, akin to the WT protein (Conicella et al. 2016; Schmidt and Rohatgi 2016; Jiang et al. 2013; Li, Chiang, et al. 2018; Schmidt, Barreau, and Rohatgi 2019; Li, Chen, et al. 2018), the transiently α-helical region may also be of key importance for LLPS of the phosphomimetic variants.

To further explore this issue, we employed circular dichroism (CD) spectroscopy which provides information about protein secondary structure. Far-UV CD spectra of WT TDP-43 LCD and 4-PM TDP-43 LCD were nearly identical (**Fig. 7A**), strongly suggesting the 4-PM variant retained the transiently α-helical region. Next, an A326P point mutation was introduced to 4-PM TDP-43 LCD to extirpate the helical structure. This helix-breaking mutation, which was shown in other studies to abolish LLPS of WT TDP-43 LCD (Conicella et al. 2016; Conicella et al. 2020), completely abrogated condensation of 4-PM TDP-43 LCD under the conditions where this phosphomimetic variant without the mutation formed liquid droplets (**Fig. 7B**). Thus, akin to the case of the WT protein, the transiently α-helical region appears to be necessary for LLPS of the phosphomimetic variant. Finally, we introduced the W334G mutation to 4-PM TDP-43 LCD; this mutation removes a tryptophan residue previously shown to be vital to WT TDP-43 LCD LLPS, while leaving the transiently α-helical structure intact (Li, Chiang, et al. 2018). W334G 4-PM protein was not able to phase separate under the tested conditions (**Fig. 7C**). Taken together, these results strongly suggested that the transiently α-helical region and key residues therein, which are crucial for LLPS of the WT protein, were still required for the phase separation of C-terminally phosphomimetic TDP-43 LCD.

**Figure 7.**
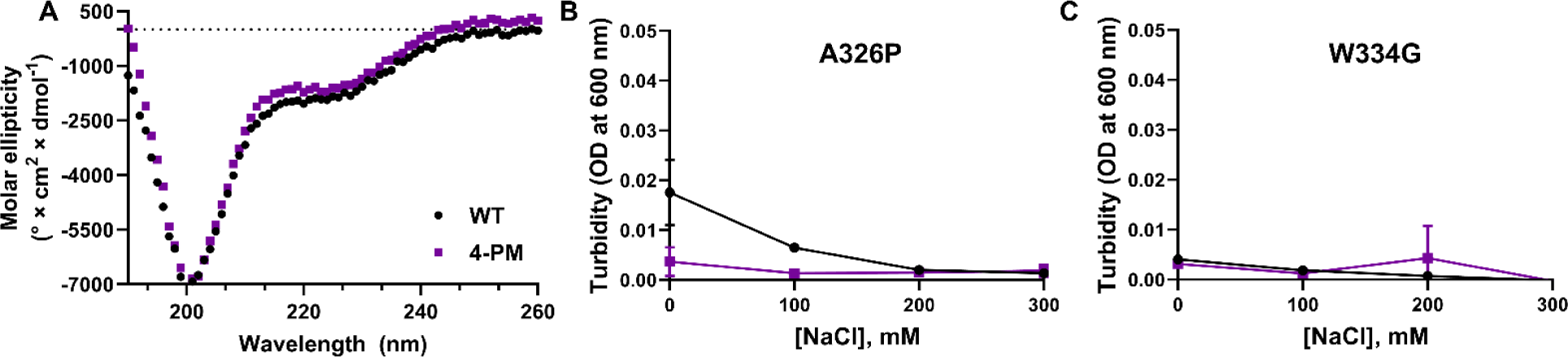
The transiently α-helical region remains critical for LLPS of phosphomimetic TDP-43 LCD variants. *A*, Far-UV circular dichroism spectra of WT and 4-PM TDP-43 LCD. *B*-C, Turbidity plots for WT and 4-PM TDP-43 LCD with the helix-breaking A326P mutation (*B*) and the aromatic-residue-removing W334G mutation (*C*). All experiments were done at room temperature, pH 7.4. OD, optical density.

## Discussion

Much of the TDP-43 within the inclusions that characterize age-related neurodegenerative diseases such as amyotrophic lateral sclerosis (ALS), frontotemporal lobar degeneration (FTLD), limbic-predominant age-related TDP-43 encephalopathy, and cerebral age-related TDP-43 with sclerosis is post-translationally modified (Neumann et al. 2006; Arai et al. 2006; Neumann et al. 2009; Nonaka et al. 2009). For example, ubiquitination and N-terminal truncation of TDP-43 have been extensively observed, with C-terminal fragments containing the low complexity domain (LCD) of the protein being a major component of amyloid aggregates in ALS and FTLD (Arai et al. 2006; Neumann et al. 2006; Nonaka et al. 2009). Phosphorylation, though, is of particular significance. Not only has phosphorylated TDP-43 been shown to specifically segregate to pathognomonic inclusions, but phosphorylation at certain C-terminal residues (e.g. S403, S404, S409 and S410) is so widespread in pathology that it is used as a biomarker of disease (Eck, Kraemer, and Liachko 2021; Hasegawa et al. 2008; Neumann et al. 2009). As such, the role of phosphorylation in the development of TDP-43 proteinopathies is an active area of investigation. One topic within this field that remains largely unexplored, however, is the effect of C-terminal phosphorylation on TDP-43 liquid-liquid phase separation (LLPS). TDP-43 forms membraneless condensates through the mechanism of LLPS, which is driven by its LCD (Conicella et al. 2016; Schmidt and Rohatgi 2016; Carey and Guo 2022; Banani et al. 2017; Shin and Brangwynne 2017; Alberti and Dormann 2019), and ever-mounting evidence has linked dysregulation of phase separation to disease (Alberti and Dormann 2019; Wang et al. 2021; Babinchak and Surewicz 2020; Zbinden et al. 2020; Haider, Boyko, and Surewicz 2023). The relative dearth of studies investigating how this potentially pathogenic process is affected by C-terminal phosphorylation, therefore, represents a crucial gap in our understanding of TDP-43 proteinopathies. To address this issue, we employed a combination of experimental and coarse-grained simulation-based approaches to ascertain the impact of these post-translational modifications on TDP-43 LCD condensation using a phosphomimetic model of the protein’s low-complexity domain.

Akin to many other LLPS-prone low complexity domains, the condensation of TDP-43 LCD is promoted by increasing the ionic strength of the buffer (Haider, Boyko, and Surewicz 2023; Martin et al. 2021; Murthy et al. 2019; Babinchak et al. 2019). This is likely due to strengthening of short-range intermolecular hydrophobic interactions as well as weakening of repulsive, long-range intermolecular electrostatic interactions (Schmidt, Barreau, and Rohatgi 2019; Babinchak et al. 2019; Babinchak et al. 2020; Li, Chen, et al. 2018). While our present study fully confirmed this salt-dependence of WT TDP-43 LCD LLPS, it also revealed that the LLPS behavior of the phosphomimetic variants studied is dramatically different. First, in the absence of NaCl, both 2-PM TDP-43 LCD and 4-PM TDP-43 LCD phase separated to a greater extent than the WT protein. These results were unexpected, as another group previously suggested (based on coarse-grained simulations only, without experimental data) that C-terminal phosphomimetic substitutions suppressed LLPS of the TDP-43 LCD (Gruijs da Silva et al. 2022). In their study, however, the simulations were run at 150 mM NaCl, where our data also showed the WT protein phase separated more robustly than the phosphomimetic proteins. Second, in sharp contrast to the WT protein, LLPS of the phosphomimetic variants exhibited a biphasic response to ionic strength, first decreasing at relatively low salt concentrations and then increasing as NaCl concentration was increased above a certain threshold level.

Interestingly, LLPS of unmodified full-length TDP-43, which has several additional, folded domains that interact with the LCD to modulate phase separation (Haider, Boyko, and Surewicz 2023), was also reported to have a biphasic salt dependence (Krainer et al. 2021), and a similar behavior was observed for full-length FUS, another RNA-binding protein associated with age-related neurodegenerative diseases (Haider, Boyko, and Surewicz 2023; Krainer et al. 2021). The non-monotonic relationship between the LLPS of these proteins and ionic strength was ascribed to attractive electrostatic intermolecular forces primarily driving phase separation in their low salt regimes, and attractive hydrophobic intermolecular forces taking over in their high salt regimes (Krainer et al. 2021). At first, this mechanism appeared congruent with the results for our TDP-43 LCD phosphomimetic variants as well, particularly given that the C-terminal phosphomimetic substitutions created polarized charge distributions in the 2-PM and 4-PM proteins that could lead to attractive electrostatic protein-protein interactions.

To test this potential mechanism, we employed a coarse-grained simulation framework. Although coarse-grained modeling does not provide the same resolution as atomistic modeling, these simulations allowed us to explore a large number of salt/protein concentration combinations on a realistic timescale. To further increase computational efficiency, we used a relatively small number of proteins (*N* ≤ 50) for each simulation in a box with periodic boundary conditions. This, however, did not appear to affect the accuracy of our results, as they showed the same trends as those generated by simulations run with five times the number of protein, albeit with minor quantitative differences (**Fig. S3**). TDP-43 LCD LLPS involves many types of interactions, including those based on π electrons (e.g., cation-π, π-π stacking, and, more recently, methionine-phenylalanine) (Schmidt, Barreau, and Rohatgi 2019; Babinchak et al. 2019; Babinchak et al. 2020; Li, Chen, et al. 2018; Mohanty et al. 2023; Li, Chiang, et al. 2018). Given the difficulties in directly including some of these interactions in coarse-grained simulations, in our framework we explicitly modeled only electrostatic interactions (via a Yukawa screened Coulombic potential) and hydrophobic interactions (via the attractive portion of a modified Lennard-Jones potential). Such simplifications notwithstanding, our simulation framework generated phase diagrams that matched all of the qualitatively important features seen in their experimental counterparts, including the monotonic and biphasic salt dependences of phase separation of WT TDP-43 LCD and the phosphomimetic variants, respectively, and the relative LLPS-propensities of the phosphomimetic variants and the WT protein at the same salt concentrations. The experimental and *in silico* data were also remarkably close at the quantitative level, with protein concentrations at the phase boundaries differing by only a few micromoles.

Our coarse-grained modeling-based analysis of forces acting on the proteins in a phase separated state did not support the possibility that attractive electrostatic intermolecular forces per se drive LLPS of the phosphomimetic variants in their low salt regimes. According to our simulations, the total intermolecular hydrophobic forces eclipsed total intermolecular electrostatic forces by two orders of magnitude in both the low and high salt regimes for all TDP-43 LCD variants studied. These *in silico* data were further supported by the observation that condensates of the phosphomimetic variants that were formed in the absence of NaCl, and thus in the absence of any salt screening, were similarly highly sensitive to 1,6-hexanediol (a disruptor of hydrophobic interactions) as condensates that were formed in the high salt regime (Kroschwald et al. 2015). In contrast, condensates of proteins whose LLPS is primarily electrostatically-driven, such as tau and C9orf72 dipeptide repeats, exhibit a considerable resistance to 1,6-hexanediol (Boyko et al. 2019; Boeynaems et al. 2017).

Altogether, our cumulative data suggest that even though LLPS of 2-PM and 4-PM TDP-43 LCD variants is driven largely by hydrophobic interactions, electrostatic forces play an important modulatory role in this reaction. This is due to polarized charge distribution in these phosphomimetic protein variants that leads to significant attractive intermolecular electrostatic interactions. We propose that these electrostatic forces, which act over longer distances compared to hydrophobic forces at low salt concentrations (Israelachvili and Pashley 1982), bring protein monomers together. This, in turn, promotes shorter-range hydrophobic interactions that are the major driving force for LLPS. Since attractive electrostatic forces decrease with an increase in ionic strength, there is an inverse salt dependence of LLPS for phosphomimetic variants in the low salt regime. In contrast, electrostatic forces experienced by WT TDP-43 LCD at low salt concentrations are repulsive. Thus, in this case NaCl promotes LLPS by simultaneously decreasing repulsive electrostatic interactions and promoting attractive hydrophobic interactions at all salt concentrations.

The main purpose of our present study was to gain mechanistic understanding of the effect of pathologically-relevant C-terminal phosphomimetic substitutions on TDP-43 LCD LLPS. To achieve this objective, it was necessary to explore LLPS over a broad range of NaCl concentrations. From a biological perspective, however, the LLPS behavior of these proteins at physiologic ionic strength is of special interest. Our data revealed that at 150 mM NaCl, WT TDP-43 LCD phase separates more robustly than the C-terminal phosphomimetic variants, even though this difference is rather modest (the *csat*’s of all three proteins fell within a 4 µmole range). It should be noted, however, that authentic phosphorylation may impart larger negative charge than a phosphomimetic substitution; thus, the difference in LLPS propensity between unmodified and phosphorylated proteins would likely be substantially larger. Furthermore, LLPS of full-length TDP-43 inside the cell is likely further modified by the N-terminal protein domains as well as other biomolecules, such as RNA and other proteins (Haider, Boyko, and Surewicz 2023; Conicella et al. 2016; Wang et al. 2018). Our present findings for homotypic TDP-43 LCD LLPS obtained using a simple phosphomimetic model provide a physicochemical foundation for the interpretation of future studies on the effect of phosphorylation on TDP-43 LLPS in a complex cellular environment.

## Materials and Methods

### Expression and Purification of WT and C-terminally phosphomimetic TDP-43

We used site-directed mutagenesis on a plasmid encoding the amino acid sequence of WT TDP-43 LCD (residues 266-414) with an N-terminal His6 tag and thrombin cleavage site (MRGSHHHHHHLVPRGS) in a pRSET-B vector to make an analogous plasmid construct for S403D/S404D (2-PM) and S403D/S404D/S409D/S410D (4-PM) TDP-43 LCD. Rosetta *Escherichia coli* was used to express the proteins, and they were purified as previously described (Babinchak et al. 2019), except that thrombin protease (Cytiva Life Sciences, Marlborough, Massachusetts) was used to cleave the N-terminal tag after FPLC and before HPLC. The post-FPLC protein was diluted by a factor of 16 into a 20 mM potassium phosphate (pH 6) solution. Immediately following this step, thrombin was added (10 U thrombin:1 mg of uncleaved TDP-43 LCD), and the solution was rocked for 48 h at ambient temperature. The cleaved protein was then concentrated (4-fold), and subsequently mixed with guanidinium hydrochloride such that the final concentration of GuHCl was 5.3 M. The mixture was then rocked for 30 minutes until the solution clarified. Subsequently, the solution was concentrated, buffer-exchanged on HPLC, and lyophilized as previously described (Babinchak et al. 2019). To determine the concentrations of protein stocks, we employed absorbance at 280 nm using an extinction coefficient of 17990 M^-1^ cm^-1^.

### Turbidity Measurements

Turbidity (absorbance at 600 nm) at 25°C was employed to measure the extent of phase separation, using a Tecan (Baldwin Park, California) Spark multimode microplate reader with Te-Cool™ active temperature control. Experiments were carried out in 20 mM potassium phosphate buffer (pH 7.4) containing NaCl at different concentrations.

### Fluorescence Microscopy Imaging

Sample was prepared in 20 mM potassium phosphate buffer (pH 7.4), at varying NaCl concentrations. Fluorescence microscopy was employed to visualize the sample, using Alexa Fluor 488-labeled protein, with a 1:20 ratio of labeled-to-unlabeled protein. 20 µL volumes of protein were plated on 35-mm dishes precoated with 1% Pluronic F-127 (Sigma Aldrich, St. Louis, Missouri). Subsequently, images were collected at ambient temperature on a Keyence (Itasca, Illinois) BZ-X710 microscope with a ×100/1.45 numerical aperture oil-immersion lens.

To prepare fluorescent probe, TDP-43 LCD was labeled with Alexa Fluor 488 NHS (succinimidyl) ester by adding 10 µL of dye in DMSO (10 mg/mL) to 100 µL of the protein (10 mg/ml) in a solution of 20 mM potassium phosphate (pH 7) buffer with 4 M guanidinium hydrochloride. The labeling reaction was conducted at ambient temperature, under constant vortex mixing. After 1 hour, excess dye was cleared via use of Zeba desalting columns (Thermo Fisher Scientific, Waltham, Massachusetts).

## Computational Methods

### Simulation Framework – Coarse Grained Parametrization

To explore the relative importance of electrostatic and hydrophobic interactions in maintaining liquid-liquid phase separation (LLPS) of TDP-43-LCD, a coarse-grained simulation framework was developed. Though coarse-grained modeling does not capture the same level of detail as atomistic modeling, its substantially lower computational expense allows for large enough timescales to observe phase separation. We implemented this framework using previously established methods (Dignon et al. 2018; Regy, Zheng, and Mittal 2021), where every amino acid was approximated as a soft sphere defined by four properties: mass (*m*), charge (*q*), hydrophobicity (*λ*), and size (*σ*) (**Table S1**) (Dignon et al. 2018). The Kapcha-Rossky Hydrophobicity scale (HPS-KR) was used for hydrophobicity values (Kapcha and Rossky 2014).

To model amino acid bonds in the simulation framework, consecutive amino acids were approximated as being connected with a “spring” and bound together with a harmonic potential of the form:

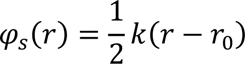

Where k is the “spring” constant between bound amino acids and *r*_0_ is the bond rest length (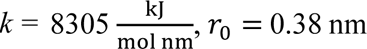 for all simulations). Two types of pairwise interactions were accounted for in our simulations: electrostatic interactions and soft interactions. Electrostatic interactions were modeled using an electrostatic potential with Debye screening.

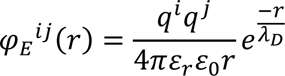

Where *q*^*i*^ and *q*^*j*^ are the charges of the interacting particles, *ɛ*_*r*_is the relative permittivity of the medium (water), *ɛ*_0_ is the permittivity of free space, *r* is the distance between interacting particles, and *λ*_*D*_ is the Debye screening length. All simulations were run in the presence of a buffer solution. Specifically, a 20 mM potassium phosphate buffer solution composed of 70% dibasic potassium phosphate and 30% monobasic potassium phosphate. The buffer solution was modeled in simulations by including the ions present therein as a contribution to the Debye screening length. (*λ*_*D*_is between 1.64 nm – 0.267 nm for tested salt concentrations). Electrostatic interactions were calculated only for particle pairs within 3.5 nm for computational efficiency. To capture soft interactions, the Ashbaugh-Hatch functional form (Ashbaugh and Hatch 2008) was used to generate a modified Lennard-Jones potential (MLJ):

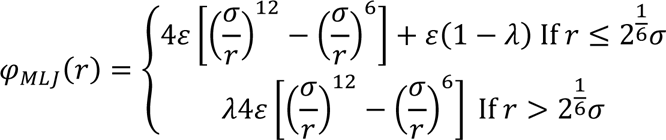

Where *ɛ* is the depth of the potential well (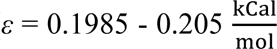 for tested salt concentrations), *σ* is the average “size” of two interacting particles, *λ* is the average hydrophobicity of two interacting particles, and *r* is the distance between interacting particles. The term

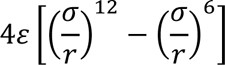

is the typical 12-6 Lennard-Jones potential. We note that the hydrophobicity parameter *λ* is present only in the attractive term of the MLJ interactions. Thus, when analyzing forces between amino acids moving forward, we define attractive MLJ interactions as hydrophobic interactions. Simulations were run in the presence of NaCl, a weakly kosmotropic ion, which means that it stabilizes hydrophobic inter-residue interactions. To account for this phenomenon in our model, we made the well depth of the MLJ potential explicitly salt-dependent with a simple first order perturbation:

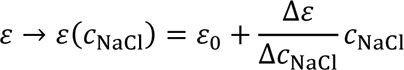

Where 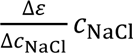 is the rate of change of well depth with respect to NaCl concentration, and *ɛ*_0_ is the baseline well depth. MLJ interactions were calculated only for particle pairs within 2 nm for computational efficiency.

### Simulation Framework - Workflow

Simulations of TDP-43 LCD and its phosphomimetic variants were performed using the Python toolkit HooMD-Blue (Anderson, Glaser, and Glotzer 2020; Howard, Panagiotopoulos, and Nikoubashman 2018). For our simulations, the number of proteins in the box was intentionally kept small (*N* ≤ 50) to allow for many regions of the protein/salt concentration parameter space to be explored. For all main text simulations, the simulation box was initially squeezed to a size of 60 nm^3^. Then, the simulation box was extended to a slab configuration (60 nm x 60 nm x 1200 nm) with periodic boundary conditions (Dignon et al. 2018; Blas et al. 2008; Silmore, Howard, and Panagiotopoulos 2017). This setup allows us to roughly mimic a larger system, though inevitably finite size effects will remain. To test the significance of finite size effects, we also ran simulations with larger numbers of proteins in a larger box (see details below). In all cases, simulations were run for 5 μs in 0.01 ps time steps propagated in time using Langevin dynamics. The system was allowed to equilibrate during the first 4 μs of the simulation, and the system was analyzed during the final 1 μs.

### Phase Separation Validation

As a proof of concept of the efficacy of the outlined simulation framework, phase separation diagrams over a range of salt concentrations (0 – 1250 mM) and protein concentrations (9.6 – 20.8 μM) were generated and compared to experimental results. The MLJ well depth and its rate of change with respect to NaCl concentration were tuned such that simulated and experimental phase separation diagrams corresponded to both 0 mM NaCl and 1250 mM NaCl for TDP-43-LCD. Thus, the experimental phase separation diagrams at 0 mM and 1250 mM salt concentration constituted boundary conditions to which the explicit salt dependence of the MLJ well depth was matched.

For each data point in the salt/protein concentration space, a simulation was run and a binary classification of “phase separated” or “dissolved” was made. This classification was rendered after visualizing the simulation results over the entire 5 μs period. Proteins with center of masses within 10 nm were classified as in the same cluster. If the number of neighbors (proteins in the same cluster) per protein was greater than 2 for the entirety of the last microsecond,the datapoint was classified as “phase separated”. Otherwise, the datapoint was classified as “dissolved”. For a given NaCl concentration, all protein concentrations greater than the lowest protein concentration at which “phase separation” was observed were assumed to be “phase separated” as well. Conversely, all protein concentrations less than the lowest protein concentration at which “phase separation” was observed were assumed to be “dissolved.” The simulated phase separation results can be seen in (**Fig. 2C**) and are shown to recapitulate the important qualitative features of the experimental diagrams.

To stress the importance of an explicitly salt-dependent MLJ well depth, the TDP-43 LCD and 4-PM phosphomimetic variant phase separation diagrams were generated with simulations run at a constant MLJ well depth 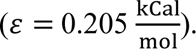. These results can be seen in (**Fig. S1**), and of note, do not recapitulate the experimental phase separation diagrams as well as the salt-dependent MLJ well depth phase diagrams do.

To validate that our approach of using a relatively small number of proteins (*N* ≤ 50) did not suffer from significant finite size effects, a subset of the phase separation diagrams was generated with protein counts between 125 and 250 for comparison. For these simulations, the simulation box was initially squeezed to a size of (60 nm x 60 nm x 300 nm), and the resulting slab was (60 nm x 60 nm x 5800 nm).

With the larger protein counts, the MLJ well depth and its rate of change with respect to NaCl concentration were recalibrated at 0 mM NaCl and 1250 mM NaCl in the same manner described previously in these methods (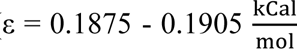 for tested salt concentrations). With these new well depth values and larger protein counts, the 0 mM, 200 mM, and 1250 mM NaCl concentration phase separation diagram columns were regenerated for TDP-43-LCD and the 4-PM phosphomimetic variant. These results can be seen in (**Fig. S3**), and of note, largely recapitulate the data produced by the smaller protein count simulations.

### Electrostatic and Hydrophobic Force Analyses

With the efficacy of the simulation framework validated, we then aimed to quantify the relative importance of electrostatic and hydrophobic interactions in maintaining LLPS over a range of salt concentrations. A natural quantity to use for this purpose is the forces generated by electrostatic interactions and hydrophobic interactions. We performed this analysis at a protein concentration of 20.8 μM, as at this protein concentration all of the TDP-43 LCD variants phase separated at every NaCl concentration tested *in silico*.

From 0 to 1000 mM NaCl, the time-averaged electrostatic and time-averaged hydrophobic forces were calculated between each pair of amino acids in the simulation box over the final microsecond of each simulation. Using the time-averaged forces, two quantities were calculated: (i) the sum of the time-averaged electrostatic forces, and (ii) the sum of the time-averaged hydrophobic forces. These results were plotted against salt concentration (**Figs. 3A and 4A**). Error bars for these data points were generated using a block bootstrapping procedure with 50 blocks and 1000 resamples

After establishing the relative importance of hydrophobic forces to electrostatic forces, we analyzed which regions of proteins were interacting most strongly in the phase separated state. To accomplish this end, the sum of the time-averaged intermolecular electrostatic forces and the sum of the time-averaged intermolecular hydrophobic forces were calculated for each amino acid residue in a pairwise association with every other amino acid residue in every protein in the box. A heat map was generated showing these sums for all residue-residue interactions (**Figs. 5, 6 and S1**).

## Code Availability

All code used for this project is available at the following Github repository: https://github.com/hincz-lab/TDP_43_Phase_Separation

## Funding

This work was supported in part by National Institutes of Health Grants RF1 AG061797 (to W. K. S.), F30 AG071339-03 (to R. H.), T32 NS077888, and T32 GM007250. The content is solely the responsibility of the authors and does not necessarily represent the official views of the National Institutes of Health.

## Declaration of Interests

The authors declare no competing interests.

## Supplementary Information

**Supplementary Figure S1.**
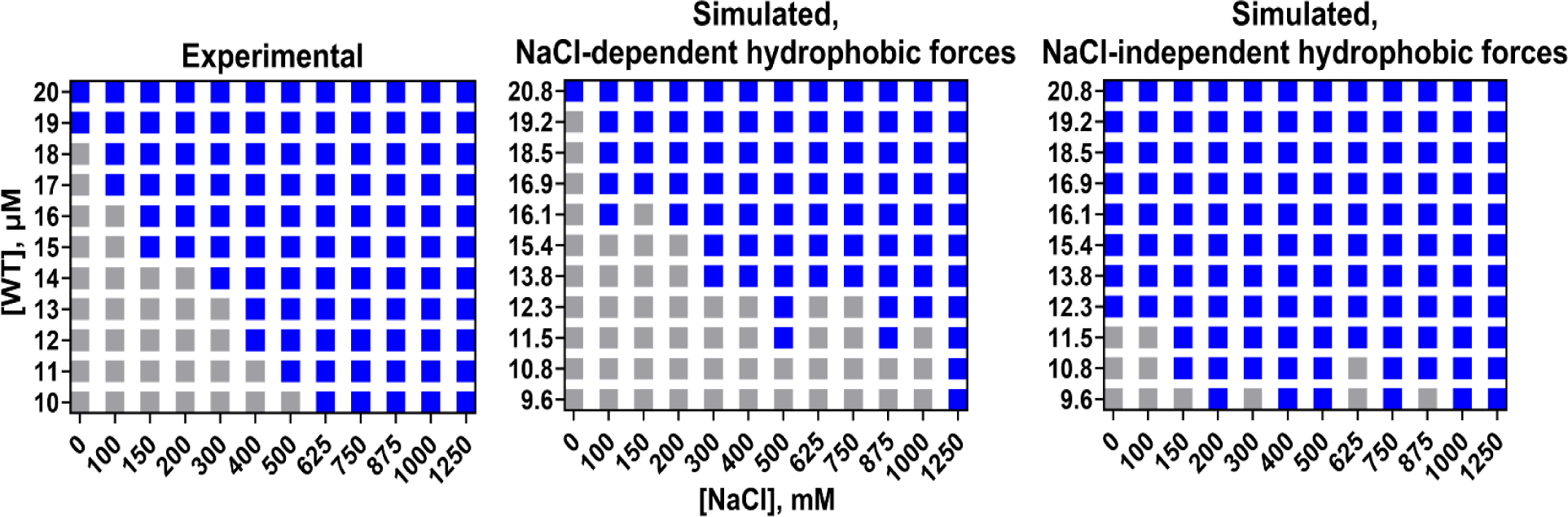
Simulations with explicit NaCl-dependent hydrophobic forces reproduce experimental trends more accurately than NaCl-independent hydrophobic forces. WT protein phase diagrams generated from experimental data (*Left*), simulations with NaCl-dependent hydrophobic forces (*Middle*), and simulations with NaCl-independent hydrophobic forces (*Right*). Experiments and simulations were done at room temperature in 20 mM potassium phosphate buffer, pH 7.4.

**Supplementary Figure S2.**
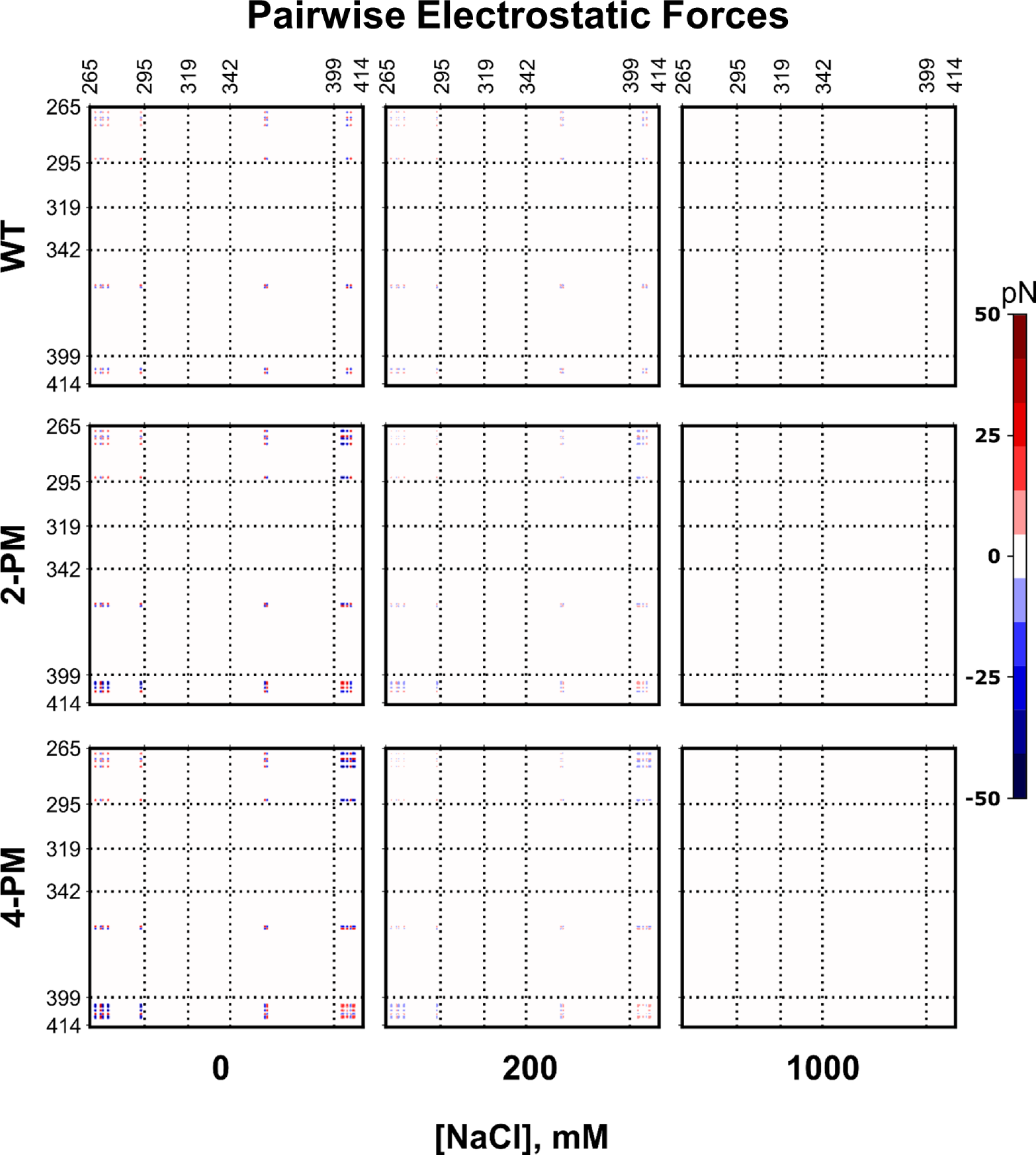
Intermolecular residue-residue pairwise electrostatic force plots of phase-separated TDP-43 LCD variants. Pairwise intermolecular electrostatic forces of WT TDP-43 LCD (*Top*), 2-PM TDP-43 LCD (*Middle*) and 4-PM TDP-43 LCD (*Bottom*) at various ionic strengths. Data were generated from 20.8 µM protein simulations.

**Supplementary Figure S3.**
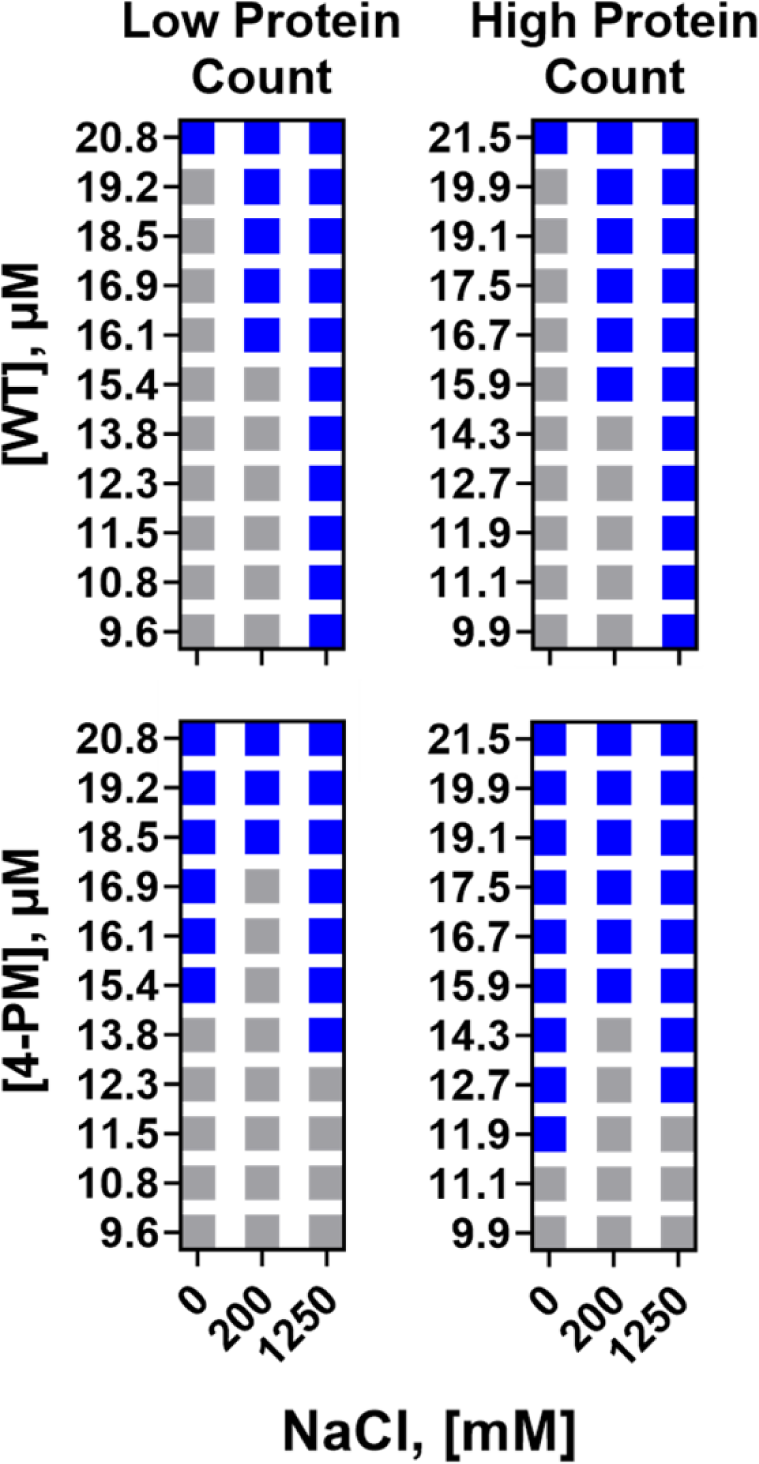
Low protein count simulation results do not substantially differ from high protein count simulation results. Phase diagrams generated from low protein count simulations (*Left*), and high (5x) protein count simulations (*Right*) for WT protein (*Top*) and 4-PM protein (*Bottom*). Simulations were done at room temperature in 20 mM potassium phosphate, pH 7.4.

**Supplementary Table S1.** Amino acid parameters for coarse-grained simulations. The fundamental properties used for simulations of each of the 20 amino acids. From Dignon et al. 2018.

| Amino acid | Mass $m$<br>[amu] | Charge $q$<br>[e] | Hydrophobicity $\lambda$ | Size $\sigma$ [Å] |
| --- | --- | --- | --- | --- |
| ALA | 71.08 | 0 | 0.730 | 5.04 |
| ARG | 156.2 | 1 | 0 | 6.56 |
| ASN | 114.1 | 0 | 0.432 | 5.68 |
| ASP | 115.1 | -1 | 0.378 | 5.58 |
| CYS | 103.1 | 0 | 0.595 | 5.48 |
| GLN | 128.1 | 0 | 0.514 | 6.02 |
| GLU | 129.1 | -1 | 0.459 | 5.92 |
| GLY | 57.05 | 0 | 0.649 | 4.50 |
| HIS | 137.1 | 0.5 | 0.514 | 6.08 |
| ILE | 113.2 | 0 | 0.973 | 6.18 |
| LEU | 113.2 | 0 | 0.973 | 6.18 |
| LYS | 128.2 | 1 | 0.514 | 6.36 |
| MET | 131.2 | 0 | 0.838 | 6.18 |
| PHE | 147.2 | 0 | 1 | 6.36 |
| PRO | 97.12 | 0 | 1 | 5.56 |
| SER | 87.08 | 0 | 0.595 | 5.18 |
| THR | 101.1 | 0 | 0.676 | 5.62 |
| TRP | 186.2 | 0 | 0.946 | 6.78 |
| TYR | 163.2 | 0 | 0.865 | 6.46 |
| VAL | 99.07 | 0 | 0.892 | 5.86 |

## Notes

### Competing Interest Statement

The authors have declared no competing interest.

